# A survey on information sources used by academic researchers to evaluate scientific instruments

**DOI:** 10.1101/253799

**Authors:** Carsten Bergenholtz, Samuel C. MacAulay, Christos Kolympiris, Inge Seim

## Abstract

Most scientific research is fueled by research equipment (instruments); typically hardware purchased to suit a particular research question. Examples range from 17^th^ century microscopes to modern particle colliders and high-throughput sequencers. Here, we studied the information sources used by academic researchers to assess scientific instruments, and reveal evidence of a worrying confluence of incentives similar to those that drove the biopharmaceutical industry to adopt controversial practices such as ghostwriting and hidden sponsorship. Our findings suggest there are little understood incentives against disclosure in the peer-reviewed literature on scientific instruments; constituting an underappreciated threat to scientific standards of trustworthiness and transparency. We believe that a public debate and subsequent editorial policy action are urgently required.

---

There is a growing concern about the reliability^1,2^, reproducibility^3,4^, and transparency of science^5-7^ and, in particular, how commercial interests might distort scientific integrity^8-10^. Studies on the biopharmaceutical industry have shown that for-profit company (‘firm’) sponsored research is more likely to reach conclusions favorable to the funding sponsor^11^, but also that such studies are considered less reliable by academic researchers^10^. In comparison, scientific instruments have profound effects on the scientific process^12-14^ and account for billions of dollars in research expenditure^13,15^, yet we know little about how the activities of firms in this industry influence the scientific process. Recent qualitative research found that firms producing scientific instruments viewed mentions of their instruments in peer-reviewed studies as valuable marketing material^16^. However, firms considered the marketing value of this endorsement substantially diminished if their employees were listed as co-authors. Even when employees made significant contributions to a paper in question, some of these firms had a policy of not being listed as co-authors. Here, we report on a study conducted to investigate whether these views reflect *bona fide* concerns.

We undertook two surveys to explore whether academic researchers devalue the information found in research co-authored by firm employees. The first survey assessed what information sources the researchers rely on when evaluating scientific instruments (survey 1: ‘importance’). The second survey measured whether co-authorship by employees of scientific instrument firms alters how reliable information in the (Materials and) Methods section of a given peer-reviewed manuscript is perceived (survey 2: ‘reliability’). See Supplementary Note 1 for further details on the surveys, respondents, and descriptive results.

## Survey 1: the importance of information on scientific instruments

With a response rate of 19%, comparable to similar studies in the social sciences (see Supplementary Note 1), we received 994 responses from U.S. and E.U. based researchers of varying academic ranks, research budgets, and academic disciplines. Our results show that input from colleagues was the main source of information (see Table S1). Peer-reviewed publications also constituted an important source, more so than scientific conferences and salespersons. We further evaluated how respondents perceived various information sources referring to scientific instruments using a 5-point Likert scale. Publications co-authored by employees from the firm producing the instrument were considered a significantly less important source than publications without firm affiliations (Table S2; *Z* = 18.26, *P* < 0.0001, Mann-Whitney *U*-test).

## Survey 2: the reliability of information on scientific instruments

A second survey of the same cohort (247 respondents, response rate of 30%; see Supplementary Note 1 for further information) provided further insights on the perception of peer-reviewed publications with co-authors from scientific instrument firms. It revealed that respondents considered information in publications co-authored by instrument firm employees less reliable (Figure 1a). In contrast to publications without any instrument firm affiliations, which ~80% deemed reliable, only 36% considered papers co-authored with someone from the instrument firm referenced in said article reliable. This pattern also applied more broadly to papers co-authored by anyone from industry, which 55% deemed reliable (Figure 1a).

**Figure 1.**
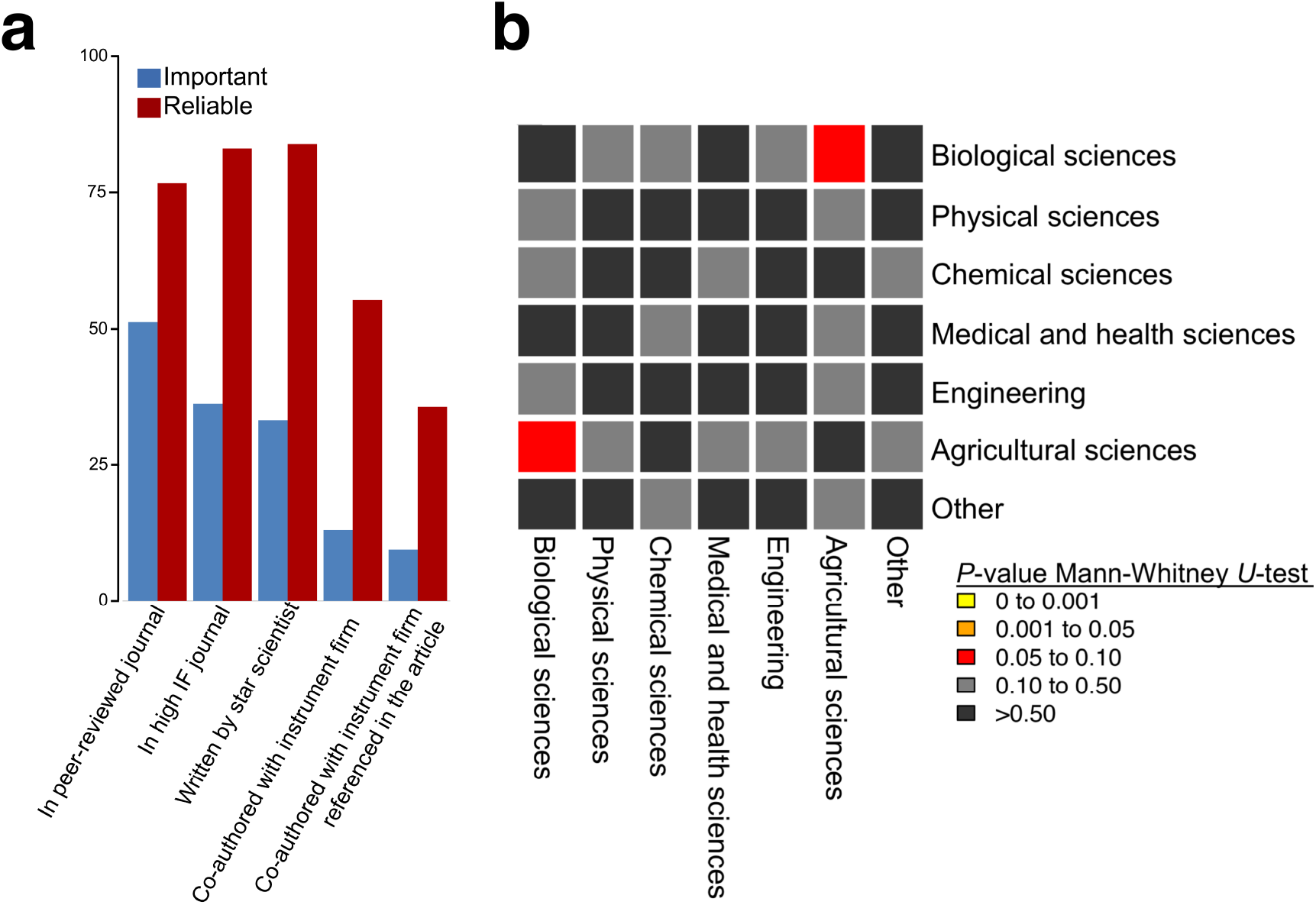
The reliability and importance of information sources on scientific instruments. (a) Illustration of how important and reliable respondents, indicated in per cent on the y-axis, consider information on scientific instruments to be in a peer-reviewed publications in general, and subcategories. (b) Heat map showing Mann–Whitney *U*-test statistics for pair-wise comparisons on the importance of firm-authored publication in different scientific fields. *P* ≤ 0.05 was considered significant. The sample number per group was ≥ 38. Please see Table S5 for details.

These data on importance and reliability were remarkably consistent across a range of potential confounding variables. These included scientific fields (Figure 1b), as well as geographic locations, degrees of entrepreneurial activity, and source of funding (see Supplementary Note 1). Interestingly, proxies of academic success (variables such as size of research budget, and the impact factor of the best journal the respondent has published in) were associated with a lower rating of the importance and reliability of publications coauthored by firm employees (see Supplementary Note 1).

## Conclusions

To our knowledge, this study provides the first systematic evidence on how academic researchers evaluate information in publications co-authored by scientific instrument firm employees. The study reveals that academics discount the importance and reliability of peer-reviewed manuscripts co-authored with scientific instrument firm employees – even when the firm’s instrument was not mentioned by the manuscript in question. The published work, thus, has reduced scientific credibility.

Descriptions of commercial scientific instruments in peer-reviewed publications have been abundant for at least half a century^17^, and researchers rely on this information source when deciding to use a given instrument^16^ (see also Table S1). However, academic researchers face a dilemma when interpreting this information source. On one hand, commercial firms have superior knowledge of their own products and are, consequently, in a position to optimize the usage of their instrument^18-20^. On the other hand, firms can have commercial incentives to misrepresent the functionalities and qualities of their instrument if it can lead to sales. Prior to this study anecdotal evidence revealed that some instrument firms prefer to hide their contributions to a scientific article^16,21^. Our data provide evidence for why: the omission of firm employees as co-authors enhances the perceived reliability of a peer-reviewed manuscript and, thus, likely sales potential of any instruments mentioned within. These dynamics are also reflected by a commercial producer of transgenic mice offering researchers monetary rewards for citations in scientific articles^22^, and scientific instruments firms promising significant discounts on instrument reagents in exchange for ‘excessive usage’ of an instrument name in scientific articles (C.B. and I.S., personal observations). This mirrors incentives for controversial practices adopted by the biopharmaceutical industry, such as ghostwriting and hidden sponsorship^23,24^. Our study provides the first systematic evidence to explain the nature of the incentives driving such behavior in the scientific instruments industry, and why, if left unchecked, it is likely to continue.

Currently the editorial impetus of peer-reviewed journals, including specialist journals such as *Nature Methods,* is to disclose any financial interests^25^. Nevertheless, there appears to be a general lack of guidelines – including by top-tier science journals (see Table S15 for a comparison) – on how and when researchers should disclose the involvement of scientific instrument firms in the production of knowledge. Such non-disclosure can leave readers unable to judge potential conflicts of interest (e.g. discounts provided on instruments) and make replication more difficult (e.g. technical assistance from a scientific instrument firm was critical for data generation, but not disclosed). Editorial guidelines have helped tackle the non-disclosure challenge in the biopharmaceutical industry^26^. As the scientific instrument industry is increasingly dominated by large corporations^27^, and expensive instruments are now commonplace in research institutes and individual laboratories^13,28^, we believe similar considerations must be applied. Without change, the existing state of affairs will continue to undermine the reliability, reproducibility, and transparency of science.

## Acknowledgements

This work was supported by a Thiess Fellowship (to S.M.), and a QUT Vice-Chancellor’s Senior Research Fellowship (to I.S.). We would like to thank Alisa Becker, Henry Sauermann, Pedro Mesquita, Angeliki Karavasili and Oana Vuculescu for their valuable input to the paper.

## Code availability

The surveys, survey data set, and associated SPSS code are available at https://github.com/sciseim/ScientificInstruments_MS.

## Author contributions

C.B and C.K. designed the research; C.B. and C.K. performed the analysis. C.B., S.M., and I. S. interpreted the results; C.B., S.M., C.K., and I.S. wrote the paper.

